# Torque and buckling in stretched intertwined double-helix DNAs

**DOI:** 10.1101/135905

**Authors:** Sumitabha Brahmachari, John F. Marko

## Abstract

We present a statistical-mechanical model for the behavior of intertwined DNAs, with a focus on their torque and extension as a function of their catenation (linking) number and applied force, as studied in magnetic tweezers experiments. Our model produces results in good agreement with available experimental data, and predicts a catenation-dependent effective twist modulus distinct from what is observed for twisted individual double-helix DNAs. We find that buckling occurs near to the point where experiments have observed a kink in the extension versus linking number, and that the subsequent “supercoiled braid” state corresponds to a proliferation of multiple small plectoneme structures. We predict a discontinuity in extension at the buckling transition corresponding to nucleation of the first plectoneme domain. We also find that buckling occurs for lower linking number at lower salt; the opposite trend is observed for supercoiled single DNAs.

## 1. INTRODUCTION

Double-helix DNA molecules are sufficiently flexible, with a thermal persistence length *A ≈*50 nm, that they often become intertwined around one another in the cell. An important example of this intertwining occurs *in vivo* at the end stages of DNA replication [1–4] due to the remnant linking of the strands of DNA in the parental double helix. Mathematicians often refer to a pair of intertwined filaments as a “2-ply” or a “2-braid” [5]; below we will describe such structures simply as braided DNAs [6, 7].

Braided DNAs have also been studied in precise singlemolecule manipulation experiments. Most such experiments have used “magnetic tweezers” to apply a constant force to a particle to which two double-helix DNAs are attached; the opposite ends of the DNAs are tethered to a surface [8–11] (Figure 1). The resulting double tether can have a controlled force applied to it using a magnetic field gradient, while at the same time, the total linking number of the two double-helix molecules can be adjusted by rotating the magnetic field so as to rotate the magnetic particle. As a result, one can study the extension of the two DNAs as a function of inter-DNA linking number (often called “catenation number”, or Ca). Experiments of this type have been used to study removal of DNA catenations by type-II topoisomerases (enzymes which change DNA topology by cutting one double helix and then passing the other double helix through the resulting gap) [9, 10, 12–14]. Braided DNAs have also been used to study the decatenation activity of type-I topoisomerases [15, 16], as well as the double-helix segment-exchange activity of site-specific DNA recombinases [17–19].

**Fig. 1.**
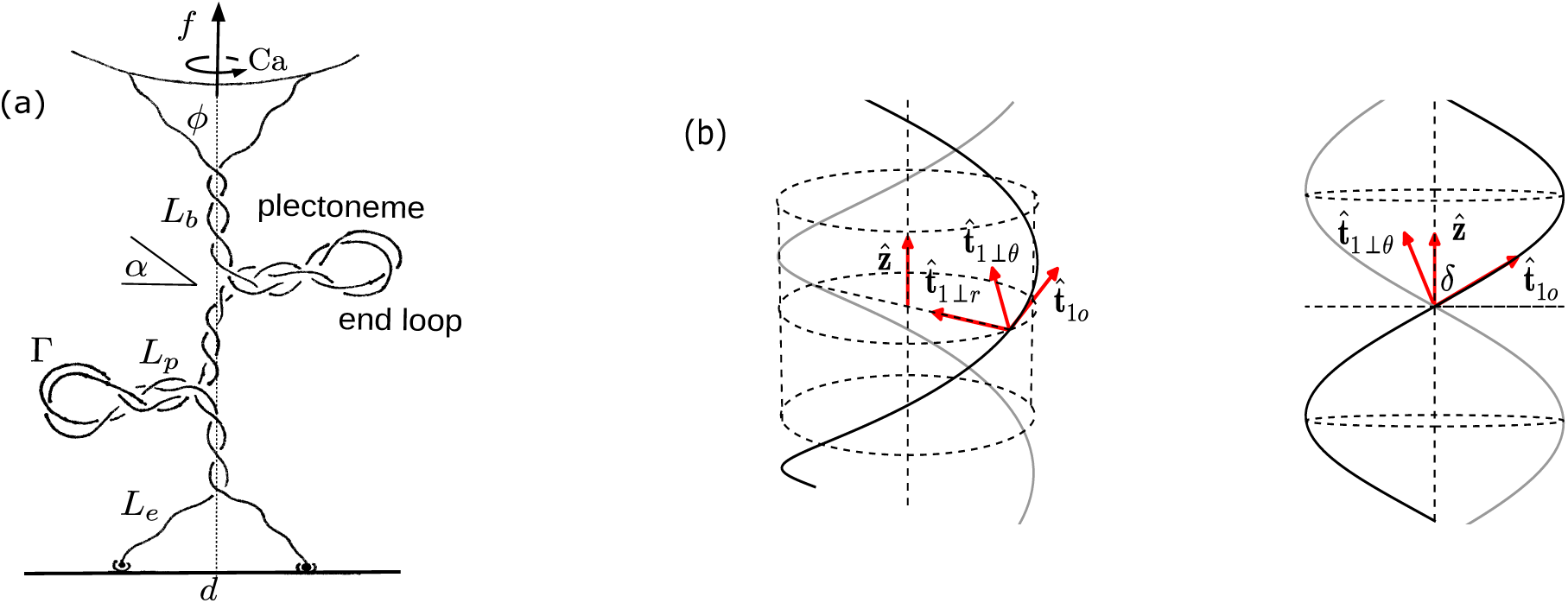
(a) Schematic of a DNA braid under torsional stress, showing coexistence of straight and plectonemically buckled states. The individual dsDNAs are able to swivel around their contact with the wall, keeping the dsDNAs from twisting. (b) Two duplex DNAs (dark and gray shaded) helically braided in a right-handed manner on the surface of a cylinder of radius *R* oriented parallel to the 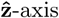, viewed from two angles. The orthonormal triad 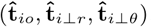, where *i* ∈ {1, 2} is shown for one of the curves. 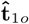 is in the direction of the tangent to the helix, 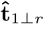 is oriented radially inward, and 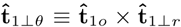. The projection of the triad on the 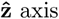 is a constant dependent on the helix parameters (Eq. 3).

Understanding these kinds of DNA-topology-changing enzyme experiments depend on the understanding of the physical properties of the DNA braids, but this has lagged behind our understanding of the simpler problem of a single twisted stretched double helix [20–28]. The reason for this is that braided DNAs are a more complex physical situation than a single supercoiled DNA; as a result, while there have been prior theoretical studies of helically intertwined DNAs [6, 11, 29–33], those works have not quantitatively analyzed the buckling (“braid supercoiling”) behavior that one would expect as catenation number is increased. While it has been assumed that the experimentally observed change in the slope of braid extension versus catenation number corresponds to the onset of braid supercoiling [9–11], the precise location and nature of braid buckling have not been theoretically understood. Two factors that make the problem of braided DNAs distinct from the mechanics of a single twisted DNA under tension are first, the lack of an intrinsic braid twist elastic modulus, and second, the dependence of the braid mechanics on the distance between the tethering points of the two double helices [9–11].

In this paper, we present the first complete theory of the mechanics of braided DNAs, treating the initial intertwining and the supercoiled-buckling behavior in a unified way. As is the usual situation in experiments [9–11], we treat the case of braids of *nicked* DNAs (double helices with a break in one of the strands, such that the broken strand can freely swivel around the intact one) to avoid the further complexity of the constraint of the double-helix linking number change on top of that of the catenation number. As a result, our model treats positive and negative catenations equivalently, a symmetry observed in experiments to a remarkable degree [9–11]. We consider various salt concentrations, DNA lengths, and intertether distances, with a focus on comparing our results to the analogous behavior of supercoiled single DNAs.

The layout of the paper is as follows. Sec. II contains detailed description of the mathematical model, where we study the braid Hamiltonian in the thermodynamic limit (Sec. II A). Free energies corresponding to a tethered braid and thermal averaging of fluctuations are discussed in Sec. II B and Sec. II C respectively. The results and predictions are contained in Sec. III, where we study braids at physiological salt (Sec. III A), as well as the effect of varying salt concentration (Sec. III B) and other finite-size effects (Sec. III C and III D).

## II. MODEL

We build a free energy model for braids considering double-helix DNAs as electrically charged semi-flexible polymers residing in an ionic solution. We define *β* = 1*/k*_*B*_*T*, and use *T* = 290 *K* for all numerical computations.

Figure 1 shows how we view braided DNA structure. The ends of two *nicked* DNA molecules are tethered to a fixed wall and a rotating bead respectively, such that the intertether distance on either end is *d*. This scenario is similar to the setup for tweezer experiments [8–11]. The beads used in experiments are large enough to safely assume no leakage of catenation number via looping of the DNA over the beads. By applying a constant force to the rotationally constrained bead it is possible to study DNA braids in a fixed force and fixed catenation ensemble.

**Fig. 4.**
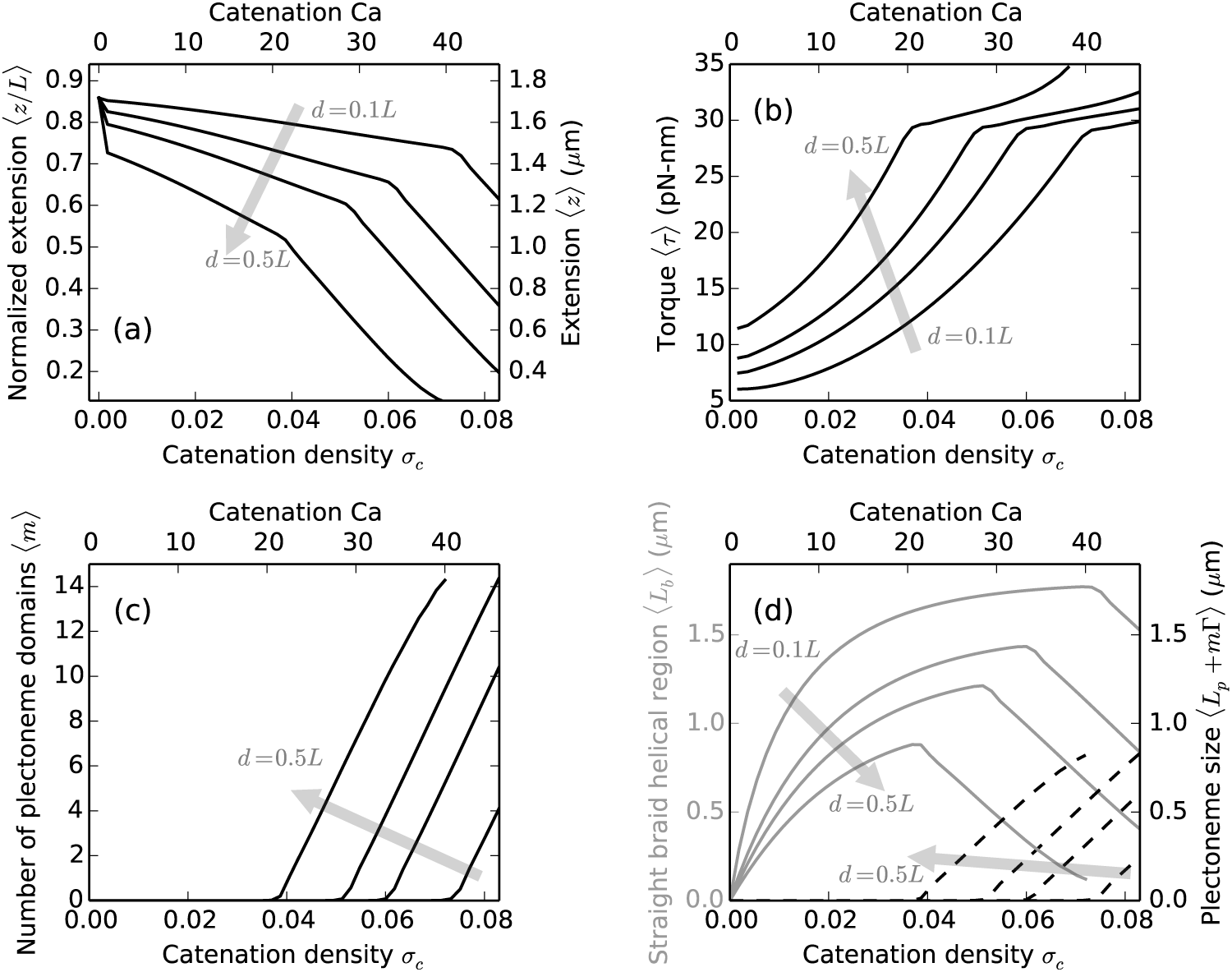
Effect of the intertether distance for braids with *≈*6 kb (*L* = 2 *μ*m) long DNAs under 2 pN force at 100 mM salt. The shaded arrows show the direction of increasing intertether distance *d* = 0.1*L,* 0.25*L, 0.35*L* and 0.5*L*, where the top and the bottom x-axes show catenation (Ca) and catenation density (*σ*_*c*_ =*Ca*/*Lk_0_) respectively. (a) Variation of relative extension (left y-axis) or extension (right y-axis) with catenation in the braid. Larger intertether distance results in a larger initial jump in the extension and lowering of the critical catenation density. (b) The torque in the braid is higher for larger intertether distances. The increase of torque per unit catenation or effective twist modulus of the braid is also higher for larger intertether distance, resulting in buckling at a lower catenation. However, the critical value of torque at which buckling occurs is a bulk property of the braid and remains roughly the same for various intertether distances, *≈*30 pN-nm at 2 pN external force. (c) The average number of plectonemes as a function of catenation showing the formation of multiple domains at all intertether distances. (d) The size of the helical straight braid (left y-axis, solid gray curves) and the total plectoneme length (right y-axis, dashed black curves) as a function of catenation. For braids with larger intertether distance, buckling occurs at relatively smaller size of the straight braid helical section due to the larger size of the triangular end regions (Figure 1).

### A The Hamiltonian

We express the Hamiltonian 𝓗 associated with two nicked double-helix DNA molecules of length *L*, held at a fixed inter-DNA linking or catenation number and under a constant applied force 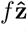as an integral over the arc length scaled by *A*, where the thermal bending persistence length of DNA *A* sets the order of magnitude of thermal deformations:

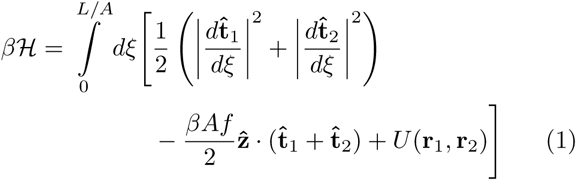

where *ξ* is the dimensionless arc length; **r**_*i*_(*ξ*) and **t**_*i*_(*ξ*)≡ =(1*/A*)(*∂***r***_*i*_/dξ*), are respectively the position vector and the tangent of the *i*-th braiding strand for *i* 1, 2. The first term in the integrand containing the sum of the squares of the local curvature of the two strands correspond to elastic bending energy of the two double helices. The second term containing the external force 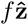 corresponds to the entropic elasticity of the two chains, and the electrostatic part of the Hamiltonian is represented by *𝒰* (**r**1, **r**2). Since we only consider nicked double-helix DNAs, there are no DNA-twist-energy terms in the above Hamiltonian, and we neglect the triangular end regions (Figure 1) till Sec. II B.

Two catenated elastic rods, many persistence lengths long, under a high stretching force form coaxial helices. We take the average shape of the braiding strands to be that of a regular helix oriented parallel to the direction of the external force 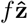(Figure 1), and propose a perturbative expansion of the braid Hamiltonian (Eq. 1) around a mean-field solution parameterized by radius *R* and pitch 2*πP* of the helix.

We expand the tangent vectors 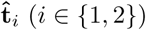 in Eq. 1 about a mean-field direction **t**_*io*_:

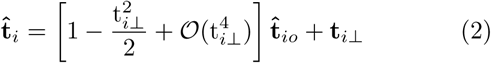

where t_*i*⊥_ = t_*i⊥r*_ + t_*⊥θ*_. We introduce two rotating right-handed orthonormal triads: 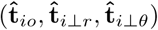, where *i* ∈ 1, 2 (Figure 1), such that the unit vector 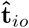points along the tangent to the mean-field helix corresponding to the *i*-th strand, 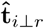 points along the radially-inward direction, and 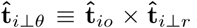. Note that the 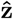 projection of the basis vectors depend only on the helix parameters:

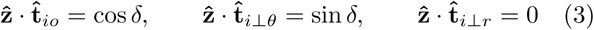

where *δ ≡* arctan(*R/P*), is the braiding angle.

The derivatives of the orthonormal basis with respect to normalized arc length *ξ* are given by the following equations:

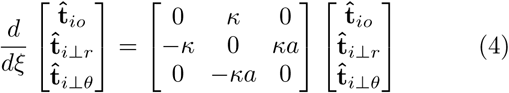

where *κ* ≡ *AR*/(*R*^2^ + *P*^2^), is the total mean-field curvature per unit persistence length of the strands and *a ≡ P/R*. The above set of equations (Eq. 4) are also known as the Frenet-Serret formulas. We neglect any space-varying component of the mean-field helix curvature [30, 34–36], which is a good approximation in the thermodynamic limit of long braids.

Using the above equations, the square of the local curvature is written as

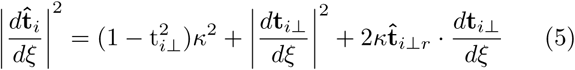

where we have neglected 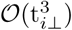 terms.

Physical micromanipulation experiments on DNA have been performed in varied concentrations of aqueous buffers, whereas *≈*100 mM Na^+^ or K^+^ is the physiologically relevant range of salt. Counterion condensation on the negatively charged DNA backbone (2*e*^−^ per base pair) results in a screened Coulomb potential over a characteristic length scale called the Debye screening length, 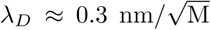 for M molar univalent salt. DNA-DNA repulsion over a few screening lengths is that of the Debye-Hückel type, *i.e.*, the electrostatic potential decays exponentially at large distances and diverges like the Coulomb potential at distances shorter than *λ*_*D*_.

The electrostatic potential due to a close proximity of two parallel DNAs has been shown to be described by Debye-Hückel interaction of uniformly charged rods [37, 38]. For helically wrapped DNA chains, also the case for plectonemes in single supercoiled DNA an empirical modification of the parallel rod potential has been shown to account for the enhancement due to helical bends in the structure [39]. Furthermore, there is an electrostatic contribution from the self interaction of the braiding helices, for which we propose an empirical form that agrees with the numerical solution of the Debye-Hückel-type self interaction (Appendix A). The total electrostatic potential energy per unit length *A* of the braid, in *k*B*T* units is given by

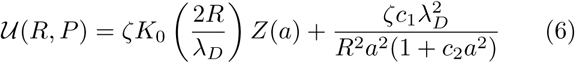

where *a* = cot *δ* and *Z*(*a*) 1 + *m*_1_*/a*^2^ + *m*_2_*/a*^4^.

The first term in Eq. 6 is the electrostatic interaction potential between the braiding strands and *Z* is the correction factor for helix curvature with *m*_1_ = 0.828 and *m*2 = 0.864 [39]. The second term corresponds to the self-electrostatic energy of a helix, where *c*_1_ = 0.042, and *c*_2_ = 0.312, are chosen to closely match the numerical solution [Figure 6]. *ζ* = 2*Aℓ*_B_*v*^2^, is the amplitude of the Debye-Hückel potential, where *ℓB* = *e*^2^/(*∊ k*_*B*_*T*), is the Bjerrum length of the solution with dielectric constant *∊* and *v* is the effective linear charge density of the double-helix DNA, which is a parameter used to satisfy the near-to-surface boundary conditions for the far-field Debye-Hückel solution [21, 25, 37–40]. We use *ℓ*_B_ = 0.7 nm corresponding to water at 290 *K*; numerical values of the effective charge *v* and the Debye screening length *λ*_*D*_, used for various salt concentrations are given in Table I. The bending persistence length of DNA *A* is also known to slightly modify on changing the salt concentration [41, 42], but we neglect such small changes as they are inconsequential to our qualitative results.

**Table I.** PDebye-Hückel parameters (the Debye length *λ*_*D*_ and the effective linear charge density *v* of the double helix [39]) and the average values of the minimized free parameters for *L* = 3.6 *μ*m, *d* = 0.42*L* and *f* = 2 pN under various salt concentrations. Comparison of the braid parameters for the straight (Ca=10) and the buckled phase (Ca=60). *R*_*s*_ and 2*π P*_*s*_ are the radius and pitch of the straight braid respectively, while *δ*_*p*_ and *α* are the braid helix angle and the superhelix angle in the plectoneme state respectively.

| Salt conc.<br>(M) | Debye length<br>$\lambda_D$ (nm) | Effective charge<br>$\nu$ ($\text{nm}^{-1}$) | Straight braid ($\text{Ca}=10$) | | Buckled braid ($\text{Ca}=60$) | | | |
| --- | --- | --- | --- | --- | --- | --- | --- | --- |
| | | | $\langle R_s \rangle$ (nm) | $\langle P_s \rangle$ (nm) | $\langle R_s \rangle$ (nm) | $\langle P_s \rangle$ (nm) | $\langle \delta_p \rangle$ ($^\circ$) | $\langle \alpha \rangle$ ($^\circ$) |
| 0.01 | 3.0 | 1.97 | 7.1 | 15.7 | 4.9 | 7.8 | 33.9 | 33.5 |
| 0.05 | 1.34 | 4.33 | 4.5 | 13.4 | 3.4 | 6.6 | 28.3 | 29.4 |
| 0.1 | 0.95 | 6.24 | 3.6 | 12.3 | 2.8 | 5.9 | 25.8 | 27.5 |
| 0.5 | 0.42 | 26.6 | 2.3 | 10.8 | 1.9 | 5.3 | 19.4 | 21.0 |

**Fig. 6.**
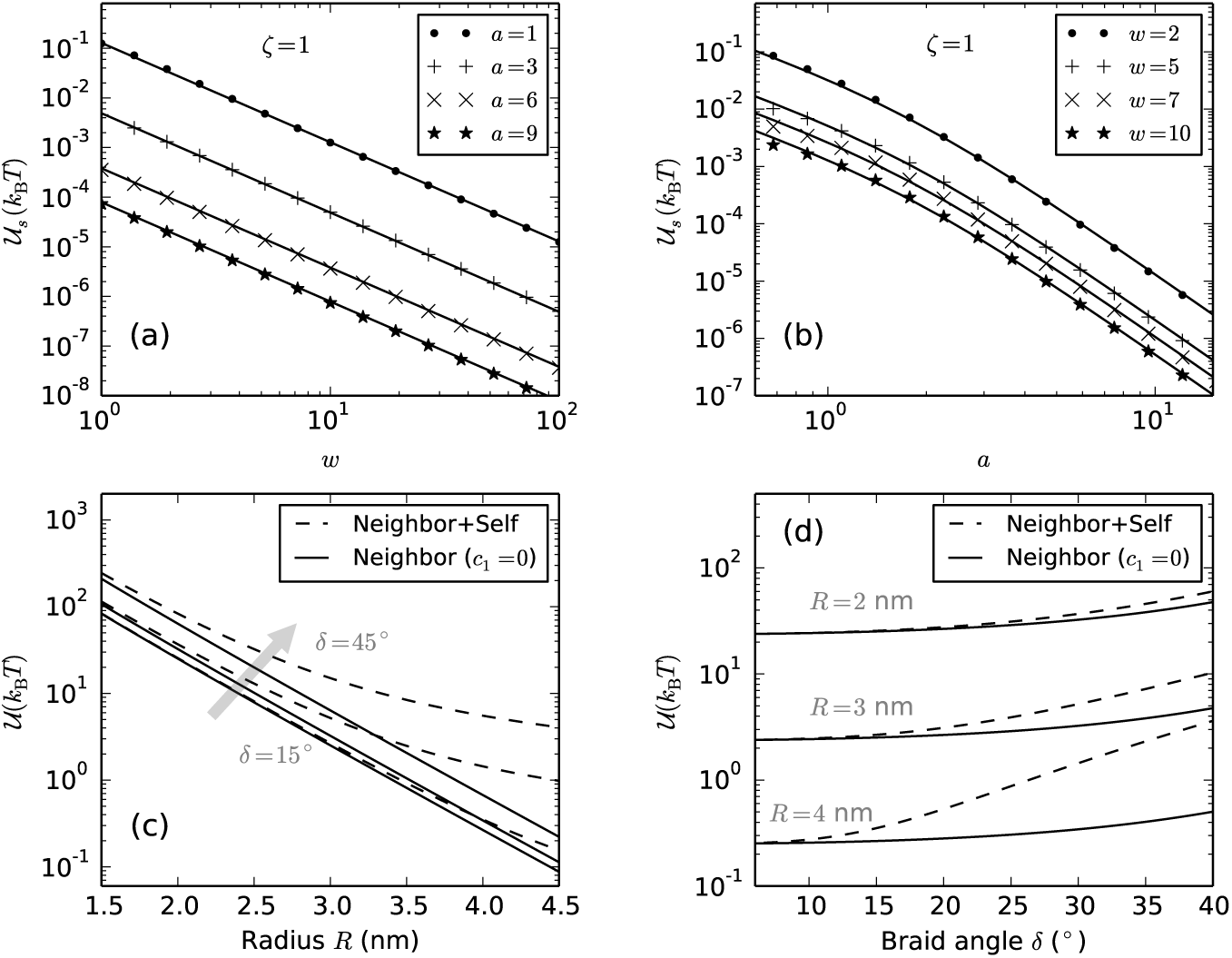
(a) Comparison of numerical evaluation (points) of *𝒰*_*s*_ (Eq. A.3) with the proposed empirical solution (Eq. A.4) (solidlines) with *ζ* = 1, *c*1 = 0.042 and *c* = 0.312, as a function of *w* for various values of *a* = 1, 3, 6 and 9. (b) *𝒰*_*s*_ versus *a* for *w* = 2, 5, 7 and 10 showing that the empirical function is a good fit to the numerical solution. (c) Plot of the total electrostatic potential *𝒰* per unit braid length *A* (Eq. A.5) with (dashed lines) and without (solid lines) the self interaction component (the second term in Eq. A.5 containing the self interaction can be set to zero by putting *c*1 = 0) versus braid radius (*R*) over a range of braiding angles *δ* = 15°, 30*°* and 45° (Table I). We used *ζ* = 2700 (corresponding to 100 mM Na^+^, Table I), *c*_2_ = 0.312 and *c*1 was chosen to be either 0 (only neighbor interaction plot, solid lines) or 0.042 (neighbor and self interaction plot, dashed line). (d) Comparison of the self and the neighbor components of the total electrostatic potential as a function of braiding angle *δ* for braid radii *R* = 2, 3 and 4 nm. The self-energy contribution is non-negligible in braids with radii ≳3 nm, which is the case ≲ 2 pN at 100 mM salt (Figure 7a).

We approximate the total braid electrostatic potential as the average potential arising from self and mutual repulsion of two coaxial helices, and consider radial fluctuations in the braid in the asymptotic limit of parallel chains. We consider small uniform deviations in the braid radius *A***w**(*ξ*) such that,

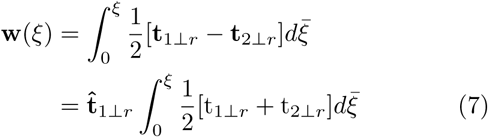

where t*i*⊥*r* are given by Eq. 2 and we assume the boundary condition **w**(0) = 0. The above definition of normalized radial deformations **w**(*?*) assumes a parallel configuration of the two strands. We define the electrostatic part of the Hamiltonian as:

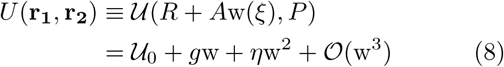

where *𝒰*_0_ ≡ *𝒰* (*R, P*), is given by Eq. 6; *g ≡ A∂U /∂R*, and *η* ≡ (*A*^2^/2)(*∂*^2^ *𝒰/∂R*^2^) is the effective modulus of the electrostatic potential. The first term gives the mean electrostatic energy per unit length *A* of braid with fixed radius and pitch, while the subsequent terms are corrections for small uniform derivation in braid radius. We neglect expansion of the electrostatic potential in the pitch of the braid because the fluctuations in the pitch are predominantly controlled by the external tension.

Now, we expand the total Hamiltonian (Eq. 1) to the quadratic order in transverse-tangent fluctuations:

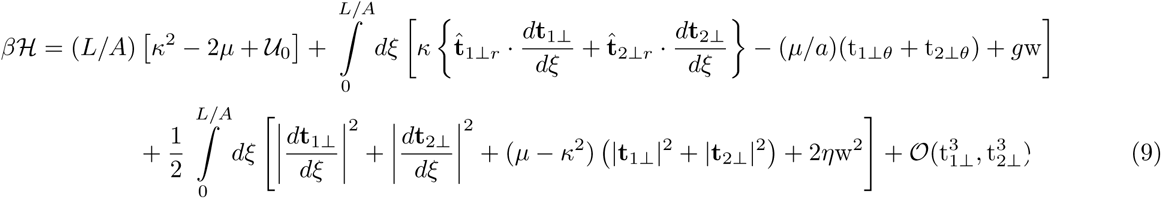

where *μ* ≡ (*β A f* cos *δ*)*/*2, is the dimensionless effective tension in each strand of the braid. The first term, associated with 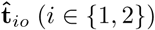 component of the tangent vectors is the leading order term that gives the total meanfield energy of the braid.

We represent the real-space components of the transverse-tangents as a sum over dimensionless Fourier modes *q*:

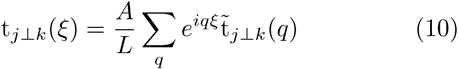

where 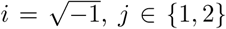, *j* ∈ 1, 2 and *k*∈ *{r, θ}*. We set the reference of the fluctuation free energy by setting the amplitude of zero-momentum transverse fluctuations to zero: 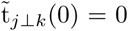, where *j* ∈{1, 2} and *k*∈ {*r, θ*}. The contribution from zero momentum is accounted for by the mean-field parameters, and this boundary condition precludes order-mean-field perturbations. Also, subject to the zero-momentum boundary condition, the second term in Eq. 9 (terms linear in t*_j_*⊥*k*) vanishes.

The third term in Eq. 9, containing quadratic transverse tangents, accounts for the free energy contribution due to Gaussian fluctuations of the two braided strands about their average helical shapes. We write the third term as a sum over the Fourier modes *q*:

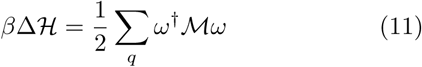

where *ω*(*q*) is a 4 *×* 1 column vector and *𝓜*(*q*) is a 4 *×* 4 Hermitian matrix such that,

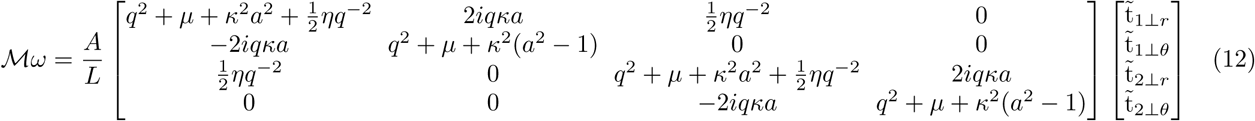

Now, we compute the fluctuation free energy in the limit of zero curvature (*κ* → 0), which simplifies the configuration to that for two fluctuating parallel chains and makes the problem analytically tractable. Also, note that in our scheme to include fluctuations in the electrostatic part of the Hamiltonian (Eq. 7) we have already assumed zero curvature.

We construct the canonical partition function for two fluctuating parallel strands:

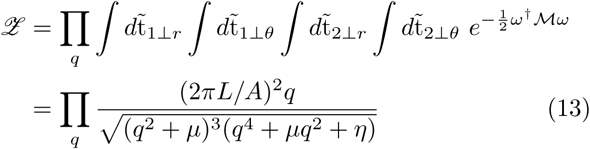

and obtain the fluctuation correction to the mean-field free energy from the partition function:

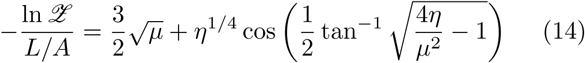

where we drop constants dependent on only the ultraviolet cutoff. The RHS of Eq. 14, which is real and positive for all positive values of *μ* and *η* gives the fluctuation free energy per unit length *A* of the braid. There are four degrees of freedom for transverse fluctuations in a stretched braid; three of them (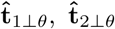, and one in 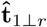where there is no relative displacement between the strands) are controlled solely by the external tension, as seen in the first term of Eq. 14. The second term accounts for fluctuations in 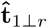that correspond to displacement of the two strands relative to one another, and is controlled by both the external tension and the electrostatic forces.

Fluctuations in the radius of the braid (*σR*) (Appendix B) decrease with increasing modulus of the electrostatic potential (*η*): *σ*_*R*_ ∼ η-^3/8^ (Eq. B.3), which implies a scaling of the fluctuation free energy with the radial fluctuations: 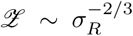. A similar scaling relation appears for the confinement entropy of a worm-like chain trapped in a rigid cylindrical tube, where the confinement entropy scales with the radius of the tube: Δ*F* ∽ 〈R〉 [43]. Theoretical studies of supercoiled DNA have used the confinement entropy scaling to account for strand undulations in a plectoneme structure [21, 27]. Again in the context of plectonemic DNA, the scaling ansatz was modified: for Gaussian fluctuations of a worm-like chain trapped in a potential well, the average radius 〈R〉 could be replaced by the radial fluctuation *σR*, which was then chosen to be the Debye length of the solution [28, 39, 44]. Indeed, we find that *σR* of the free energy minimized braid is of the order of the Debye length (Figure 7b, Table I). The existing literature on plectonemic and braided DNAs, to the best of our knowledge uses the confinement entropy scaling approach to account for strand undulations [21, 27–33, 39]. Our calculations treat fluctuations systematically and without a scaling ansatz, and produce the previously assumed scaling behavior.

**Fig. 7.**
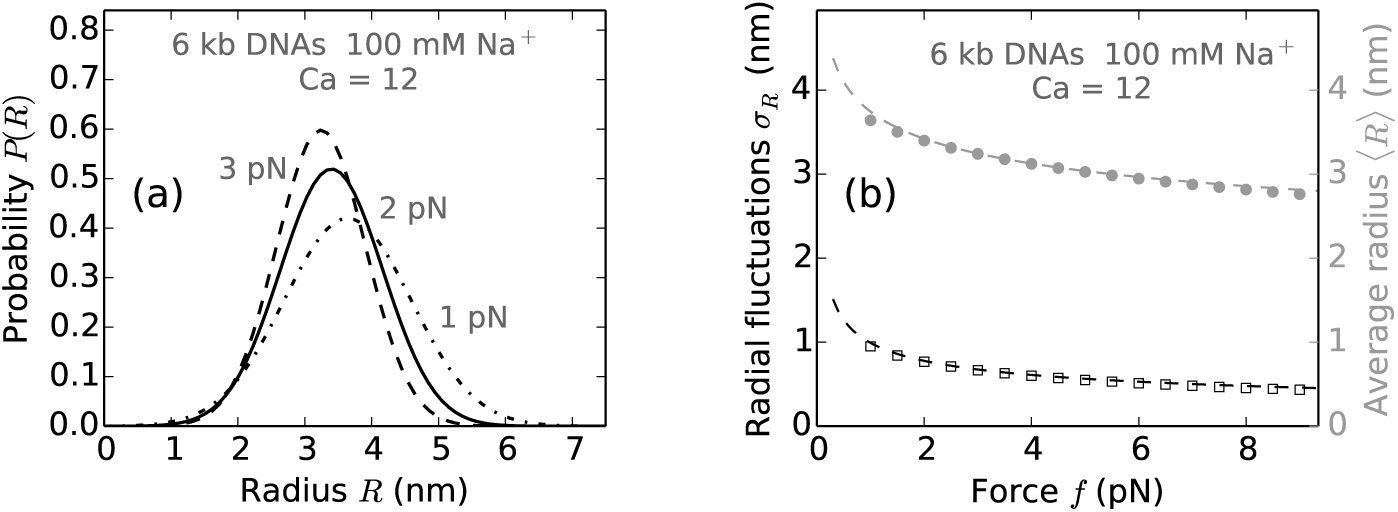
Radial fluctuations in braid with *L* = 2 *μ*m, *d* = 800nm and Ca= 12. (a) Probability distribution of braid radii (Eq. B.4), plotted for applied forces of 1 pN (dot-dashed line), 2 pN (solid line) and 3 pN (dashed line). Note the broadening of the distribution at lower forces indicating higher fluctuations. *R* = 1 nm corresponds to the excluded volume radius of the DNA molecules, included while calculating the average radius via free energy minimization but not explicitly taken into account in plotting Eq. B.4. (b) Variation of the standard deviation of the radial distribution *σ*_*R*_ (black open squares) and the average value of the radius *〈R〉* (shaded filled circles) with the external tension (*f*) on the braid. The dashed lines correspond to best-fit equations: *σ*_*R*_ = *σ*_0_(*βAf*)^−0.35^ and 〈R〉 = *R*_0_(*βAf*)^−0.13^, where *σ*_0_ = 2.4 nm and *R*0 = 5.2 nm. The exact power laws depend on the catenation in the braid, however, both the average value and the fluctuations in braid radius always decrease with increasing force.

### B. Mean-field theory

In this section, we develop the free energy expressions for a tethered braid (Figure 1), where the total length of each of the braiding molecules is partitioned into a force-extended state (straight phase), and a plectonemically buckled state (plectoneme phase). The plectoneme state also consists of a braid “end loop”, a teardrop-shaped loop at the end of every plectoneme structure (Figure 1).

The amount of catenation per helical repeat of the DNA molecules is defined as the catenation density in braids, *sc =* Ca*/*Lk_0_ (Lk_0_ = *L/h*, where *L* is the contour length of each DNA and *h* = 3.6 nm, is the length of one helical repeat of double-helix DNA). Total catenation (Ca) is divided between the straight phase (Ca_*s*_) and the plectoneme phase (Ca_*p*_), which are further redistributed between twist and writhe as dictated by minimization of the total free energy.

#### 1. Straight braid

The length of each double helix in the straight phase *L_s_* is divided into two parts: (1) the helical intermolecular wrappings of length *L_b_*, such that 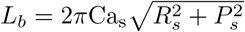, where *R_s_* and 2*P_πs_* are respectively the radius and the pitch of the helical interwounds; and (2) the end regions of length *Le* (*Le* = *Ls -Lb*), which do not contain any inter-molecular links and connect the helical wrappings to the tethered points (Figure 1). The mean-field energy of the straight braid is obtained using the leading order term in the expansion of the Hamiltonian (the first term in Eq. 9):

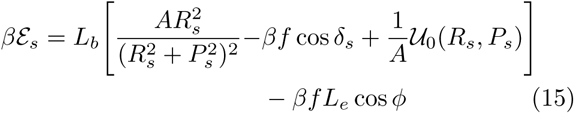

The first term (with the brackets) corresponds to the helical region of the straight phase, which is a sum of free energy contributions from elastic bending, force-extension, and electrostatic repulsion respectively. Here *δ*_*s*_ is the braiding angle (tan *δ*_s_ = *R*_*s*_/*P*_*s*_) in the straight phase. The second term contains the force-extension free energy of the end regions, where *ϕ*is the opening angle at the end of the braid (sin *ϕ*= *d/Le*, where *d* is the intertether distance, see Figure 1). Note that for a given length (*Ls*) and catenation (Cas), the radius (*Rs*) and the pitch (2*π*_*s*_) of the braid are the only free parameters in the free energy of the straight phase, minimizing which we obtain the equilibrium state (Table I).

The fluctuation correction to the mean-field free energy in the straight phase is obtained from Eq. 14,

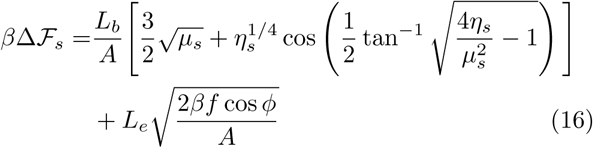

The first term (with the brackets) corresponds to the fluctuation contribution to the free energy of the helically wrapped section of the straight phase, where *μ*_*s*_ = *μ*(*R*_*s*_, *P*_*s*_) and *η*_*s*_= *η*((*R*_*s*_, *P*_*s*_) (Eq. 8 and 9). The second term corresponds to the worm-like-chain fluctuations of the end regions and is obtained by plugging *μ* = (*βAf* cos *φ*)*/*2 and *η* = 0 in Eq. 13.

#### 2. Plectonemically buckled braid

The plectonemic braid is a buckled structure where the braid centerline writhes around itself (Figure 1). Buckling achieves a lower energy state by the release of torque in the braid, due to increase in the writhe contribution to the total linking number. Total writhe in the plectoneme formed of superhelical wrappings of the braid with total double helix length *L*_*p*_, superhelix opening angle *α*, and superhelical radius *ℛ*_*p*_ is given by

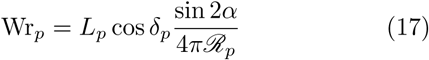

where *δ*_*p*_ is the helix angle of the braid in the plectoneme [21, 39, 45].

As mentioned above, every plectoneme domain is accompanied by a finite sized loop-shaped structure where the braid bends back (Figure 1). The braid end loop presents an energy cost to nucleation of a plectoneme domain; thermodynamically, the situation is similar to plectoneme nucleation in supercoiled single DNAs [26, 27, 46, 47]. The equilibrium size of the end loops Γ is obtained by separately minimizing the elastic energy cost of forming them:

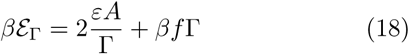

where the first term is the bending energy contribution, and the second term is the work done in decoupling the plectoneme end loop from the external tension *f* [26, 27]. We use *ε* = 16, corresponding to a “teardrop” geometry of the loop [48–50]. Minimizing *ε*_Γ_ (Eq. 18) yields,

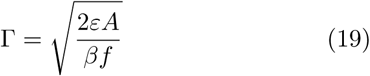

Considering the writhe contribution of an end loop to be unity (W_rΓ_ *≈*1), the total catenation in the plectoneme phase (Ca_*p*_) made up of *m* domains is partitioned into the twist (Tw_*p*_), containing the local twisting of the braid, and the writhe (Wr_*p*_), reflecting the overall structure of the plectoneme.

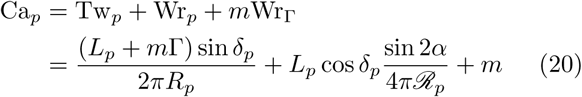

where *R*_*p*_ is the radius of the braid. Considering a simple geometric picture of closely-packed braids, we set the superhelical radius to be twice the braid radius in the plectoneme: *ℛ*_*p*_ = 2*R*_*p*_. As a simplifying assumption, we ignore local structural rearrangements in the plectoneme that may lead to spatially-varying mean-field superhelical radii.

The mean-field free energy of the plectoneme phase is given by

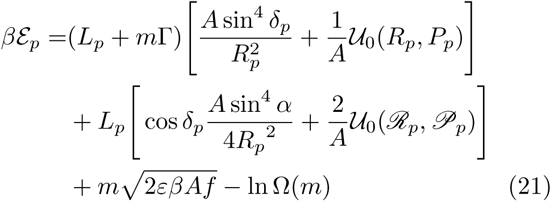

where the first bracketed term is the sum total of elastic bending energy and electrostatic energy of the braid inside the plectoneme, obtained from the mean-field term in Eq. 9. The second bracketed term is the energy contribution from elastic bending and electrostatic repulsion in the superhelix, where 2*π*P_*p*_ and 2*π*𝓅_*p*_ are the braid pitch and the superhelical pitch in the plectoneme respectively. The factor of 2 multiplying the superhelix electrostatic term is because the length of the superhelix is half of that of the DNA length in the plectoneme while the effective charge is two times that of the double helix. The third term corresponds to the elastic energy of *m* braid end loops (Eq. 18), and finally, the logarithm term is the free energy associated with the configuration entropy of *m* loops. The origin of this entropy is from the following two sources: (1) sliding of a plectoneme domain along the braid contour, and (2) exchange of DNA length among the plectonemic domains. We define *v* the arc length corresponding to unit twist in the braid 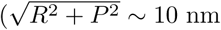 see Table I) as the characteristic length distinguishing these energetically degenerate but structurally distinct states. The total number of such states (Ω) for a plectoneme phase constituted of *m* domains (where *m* ≥ 1) can be written as a product of two combinatorial factors [27]:

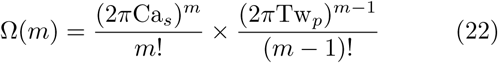

where the first term corresponds to the sliding entropy of *m* loops (2*π*Ca_*s*_ is the total number of possible plectoneme nucleation sites) and the second term is the number of distinct configurations associated with the exchange of DNA length among the domains. Note, for a plectoneme of given length (*L*_*p*_), catenation (Ca_*p*_) and number of domains (*m*), the total free energy has two free parameters (namely, *δ*_*p*_, *α* and *R*_*p*_ constrained by Eq. 20) that determine the equilibrium structure (Table I).

Similar to the straight braid case (Eq. 16), the fluctuation free energy correction to the mean-field energy of the braid in the plectoneme is obtained from Eq. 14 using *μ*_*p*_ = *μ*(*δ*_p_) and *η*_p_ = *η*(*δ*_*p*_, α):

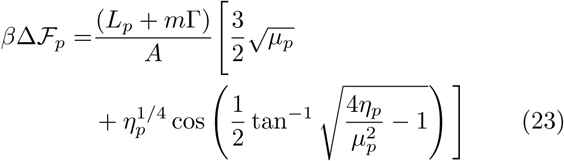

The above expression gives the total free energy associated with worm-like-chain fluctuations in the braiding strands forming the plectoneme structure.

### C Thermal averaging: partition function

In an ensemble of fixed catenation and fixed force, the total free energy of the braid can be obtained by minimizing the sum total of the straight and the plectoneme phase energy. We thermally average over states with all possible plectoneme lengths and number of domains, where the free energy in each state is minimized with respect to the partition of the total linking number, thus ensuring torque balance between the two structural phases. The free energy of the braid for each fixed *Lp* and *m* is obtained via numerical minimization over Ca*s* (Eq. 15, 16, 21 and 23),

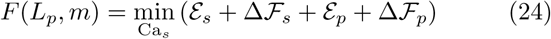

The above minimization is constrained by conservation of total catenation (Ca = Ca_*s*_ + Ca_*p*_) and total DNA length (*L* = *L*_*s*_ + *L*_p_ + *m*Γ). The states described by Eq. 24 for all possible values of *L*_*p*_ and *m* are then thermally averaged over to construct a partition function:

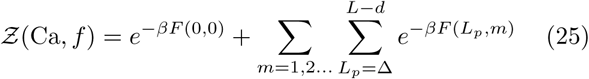

where the first term is the purely straight phase, and the second term corresponds to a sum over all possible coexistence states. The sum over *L*_*p*_ in Eq. 25 is done numerically using a Δ = 1 nm mesh. An averaging scheme as described above takes into account various thermally accessible equilibrium states which allow the possibility of torque fluctuations in the fixed catenation ensemble. A similar approach was taken in Ref. [27] to study single supercoiled DNA.

Equilibrium values of the end-to-end distance (*z*), the torque in the braid (*τ*), and the size of the helical wrappings in the straight phase (*L*_*b*_) are obtained from the partition function (Eq. 25):

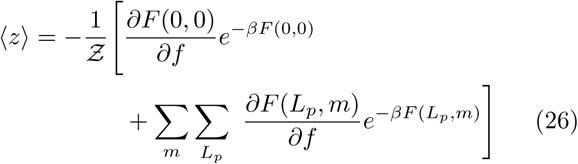

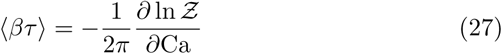

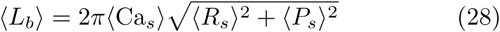

Now, the average values of the pure-state free variables *X* and the coexistence-state free variables *Y* are computed from Eq. 25 as follows,

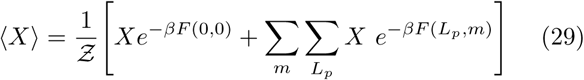

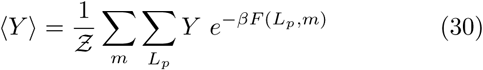

where *X ∈ {Rs, Ps,* Ca*s}* and *Y ∈ {dp, a, Lp, m}*.

## 3. RESULTS

### A. Braids at 100 mM salt

Figure 2a shows the comparison of theoreticallypredicted extension curves for various forces: 1.25 (lowest curve), 2, 3 and 4 pN (highest curve) with experimental observation for 2 pN (filled circles) at 100 mM univalent salt concentration [11]. The size of the intertether distance *d* being comparable to the length of the braiding molecules results in a sharp decrease in extension when the first catenation is added. The decrease is due to the formation of the first helical bend in the braid along with the end-regions from the zero-catenation parallel configuration. The extension shortening is used to estimate the intertether distance by simply using the Pythagorean theorem [9–11]. Notably, the intertether distance *d* is a parameter that has not been controlled in experiments to date.

**Fig. 2.**
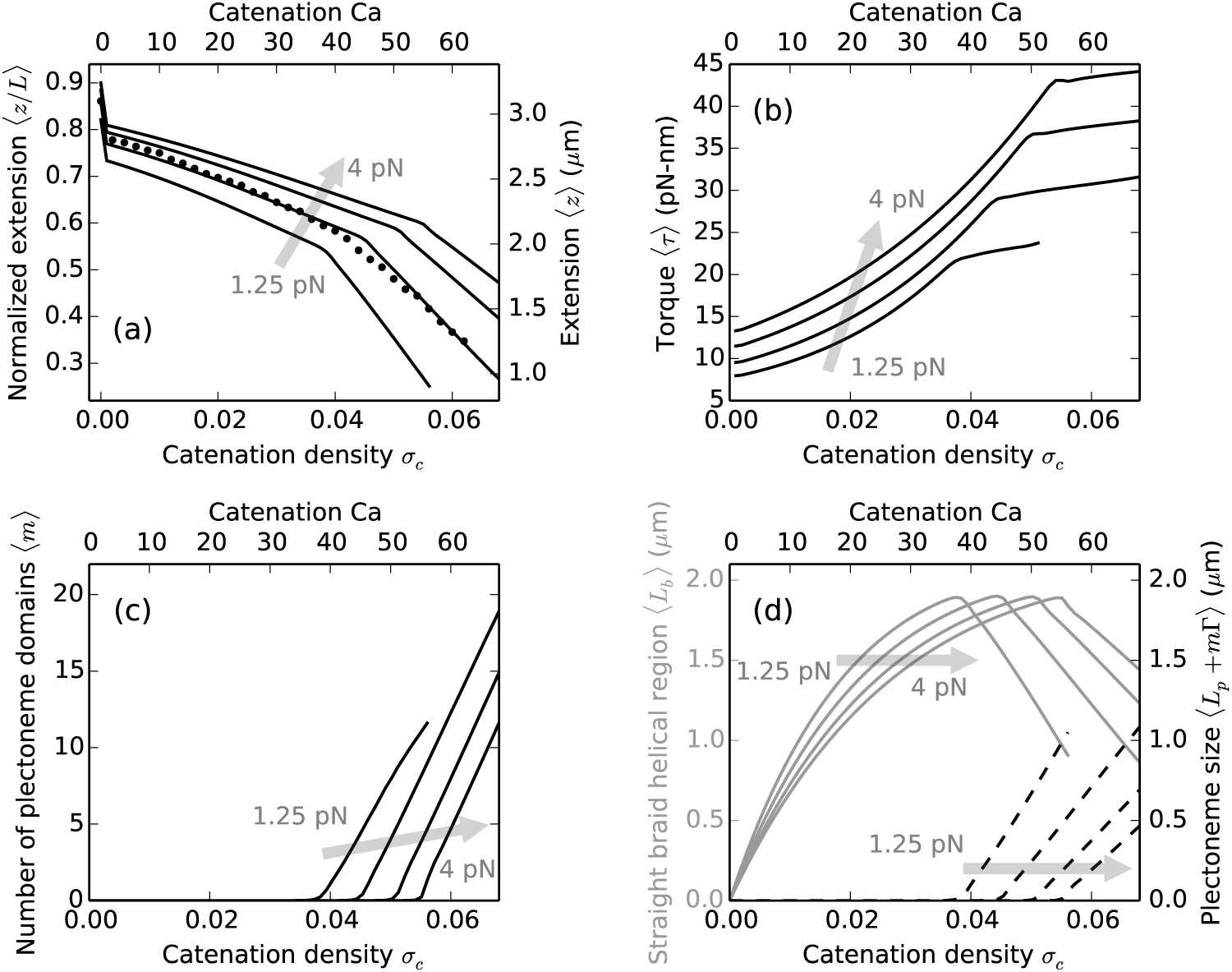
FIG. 2. DNA braids at 100 mM monovalent salt under various forces, the shaded arrows show the direction of increasing force. Theoretical predictions are for *≈*11 kb (*L* = 3.6 *μ*m) long double helices, tethered 1.5 *μ*m (*d* = 0.42*L*) apart. Catenation (Ca) and catenation density (*σc* = Ca*/*Lk0) are plotted on the top and the bottom x-axes respectively. (a) Relative end-to-end distance (left y-axis) or extension (right y-axis) versus catenation. Lines are theoretical predictions for 1.25 (lowest curve), 2, 3 and 4 pN (highest curve) force, while filled circles are experimental data at 2 pN [11]. The change in slope of the lines corresponds to plectonemic buckling transition, which is at a higher catenation for larger external tension due to increased stability of the force-coupled straight state. The kink at the onset of buckling transition is related to the plectoneme-nucleation cost presented by the braid end loop. (b) Torque in the braid shows a non-linear increase in the straight phase, and continues to increase in the coexistence phase but with a much weaker slope. The torsional stress is released in the coexistence phase due to the contribution from plectoneme writhe (Eq. 17). (c) Number of plectonemic domains versus catenation, showing that the buckled phase is characterized by multiple plectoneme domains. Nucleation of new domains causes the increase of torque in the coexistence phase, as opposed to a constant torque expected in the case of a single plectoneme domain. (d) Plot of the size of the straight-phase helical region 〈*L*_*b*_〉 (left y-axis, solid gray curves) and the size of the plectoneme region *Lp* + *m*G (right y-axis, dashed black curves) as a function of catenation. *L*_*b*_ increases in the straight phase with catenation till the buckling point, after which 〈*L*_*b*_〉 decreases as DNA length is passed into the plectoneme phase, also seen in the increase in the total size of the plectoneme.

Further addition of catenation decreases the end-to-end extension of the braid (Figure 2a) due to double helix length being passed from the end-regions to the helically wrapped section. The size of the helically-wrapped straight braid increases with catenation and reaches a maximum just before the onset of buckling (Figure 2d). Elastic bends in the braiding double helices generate torsional stress, which increases non-linearly with catenation (Figure 2b); this nonlinearity has been seen in previous models of the straight braid [11, 29, 30]. When the torque reaches a critical value, which mainly depends on thermodynamic parameters such as the external force, nucleation of the first plectonemically buckled domain becomes energetically favorable.

The onset of buckling can be identified as a *knee* in the extension plots (Figure 2a), past which DNA length is passed into the force-decoupled buckled phase (Eq.21), resulting in a steeper decrease of the end-to-end extension. The torque in the braid shows a small nonmonotonic “overshoot” at the buckling transition, and continues to increase with a small slope in the coexistence phase. In the coexistence region, the writhe contribution to the total linking number reduces torsional strain in the braid. The abruptness of the buckling transition owes to the finite-energy cost to nucleating a plectoneme domain, vis-`a-vis the plectoneme end loop. The coexistence phase at 100 mM salt is characterized by multiple domains of plectoneme (Figure 2c), where the number of domains is equal to the equilibrium number of plectoneme end loops. The total size of the plectoneme phase increases after the buckling transition (Figure 2d), where DNA length is transferred to the buckled region from the straight phase.

Higher external tension lowers the total energy of the straight braid (Eq. 15 and 16), resulting in a higher end-to-end extension at a given catenation (Figure 2a). The stabilization of the straight phase upon increasing force has an effect of delaying the buckling transition, *i.e,* buckling occurs at a higher value of catenation. The torque is higher in both the straight and the buckled braid under larger external tension (Figure 2b).

Stretched, supercoiled single DNAs at 100 mM salt also shows a buckling transition separatinga force-extended phase from a plectoneme-coexistence phase [21, 42, 51], although the mechanical response of braids is fundamentally different than that of supercoiled DNA. The torque in a supercoiled single DNA increases linearly with the linking number [46, 52], as opposed to a non-linear increase in braids (Figure 2b). The linearity of torque in supercoiled single DNA arises from a constant effective twist modulus in the double helix (*C ∼ ∂τ /∂σ_sc_ ≈* 100 nm [22, 42, 46]), which is attributed to the strong basepairing interactions holding the DNA strands together. Conversely, braids are soft structures (the two braiding molecules are not attached to each other), where twist-stiffening occurs as the catenation is increased, making the twist modulus of braids a quantity that depends on catenation as well as external parameters like the salt concentration (Sec. IIIB).

### B. Effect of salt concentration

Lowering the ionic strength of the solution causes an increase in excluded volume of the double-helix DNA in the solution, due to less screening of the negative charge on the double-helix backbone. At low salt, the Debye length (*λ*_*D*_) of the solution increases, causing the Coulomb-repulsion effect to propagate a longer distance before it is cut off, thus increasing the effective diameter of the double helix.

**Fig. 3.**
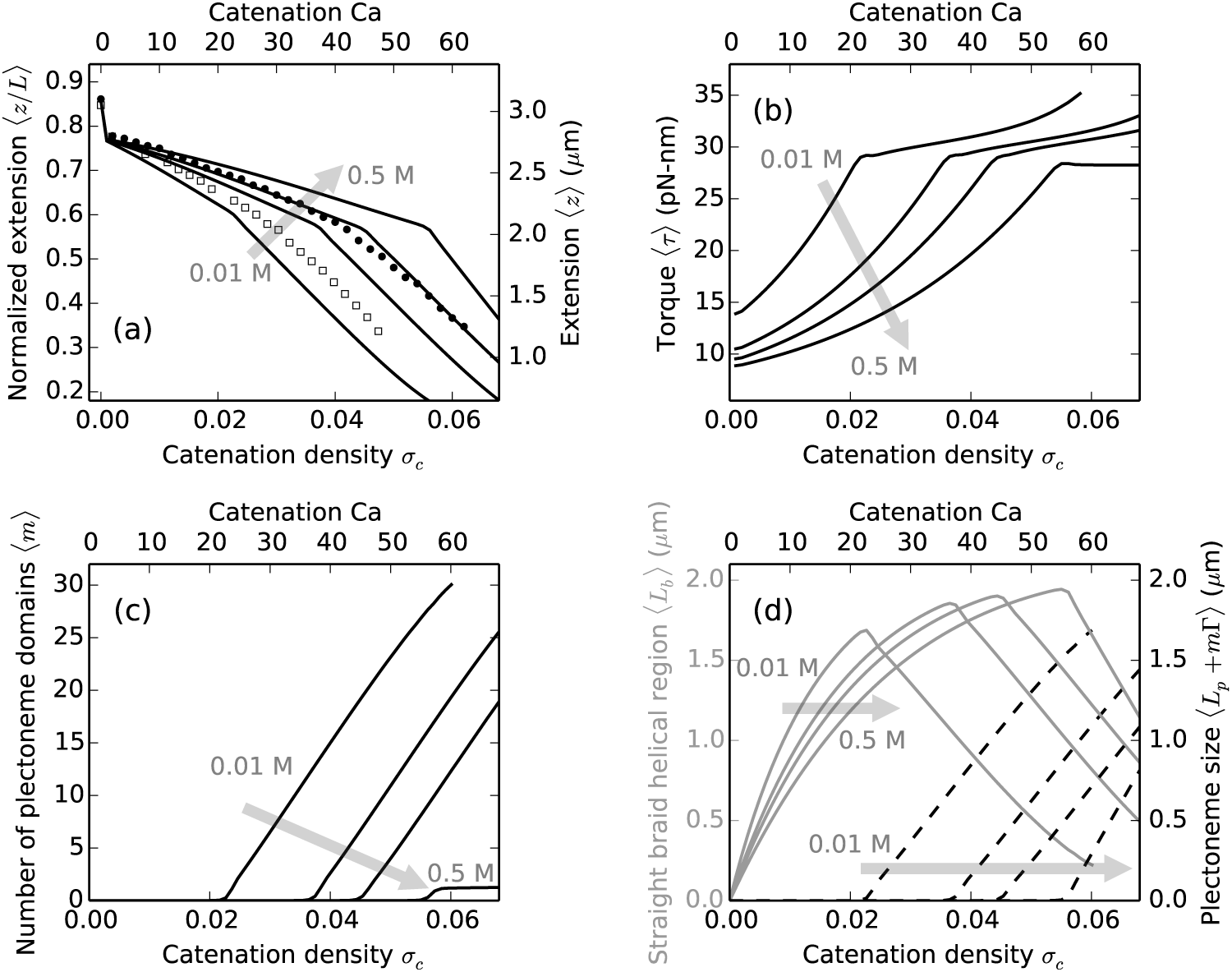
Effect of salt concentration on *≈*11 kb (*L* = 3.6 *μ*m) DNA braids under 2 pN force. The tether points are 1.5 *μ*m (*d* = 0.42*L*) apart. Theoretical curves are plotted for (0.01, 0.05, 0.1 and 0.5) M salt concentrations, where the shaded arrows show the direction of increasing salt concentration. Catenation (Ca) and catenation density (*σ*_*c*_ ≡ Ca/Lk_0_) are plotted on the top and the bottom x-axes respectively. (a) Relative end-to-end extension (left y-axis) or extension (right y-axis) versus catenation shows smaller extension and buckling at a lower catenation for lower salt concentrations. Low salt increases the effective DNA diameter, which effectively increases the twist elasticity of the braid, thereby decreasing the stability of the straight phase. The filled circles (0.1 M) and open squares (0.01 M) are experimental data reproduced from Ref. [11]. (b) Torque in the braid shows a non-linear increase in all salt conditions. Also, twist stiffening occurs faster for braids at lower salt concentration due to the larger radius of the braid. The critical buckling torque, being a thermodynamic variable does not vary significantly with the salt concentration. (c) The number of plectoneme domains versus catenation or catenation density, showing nucleation of multiple domains of plectoneme at lower salt concentrations, while a single plectoneme state is favored at higher salts. Smaller excluded diameter of the braid at higher salt makes the superhelical bending in the plectoneme phase favorable over nucleation of new domains. (d) The size of the straight phase helical wrappings 〈*L*_*b*_〉 (left y-axis, solid gray curves) and the size of the plectonemic phase 〈*L*_*p*_ + *m*Γ〉 (right y-axis, dashed black lines) versus catenation. *L*_*b*_ increases faster for lower salt concentrations due to larger braid radii (Table I).

Figure 3a shows braid extension curves under various salt concentrations (10, 50, 100 and 500) mM at 2 pN force. The buckling transition occurs at a lower catenation for lower salt concentrations. The larger effective DNA diameter increases the braid radius at lower salt conditions (Table I), which destabilizes the straight phase and consequently causes buckling at a lower catenation number. The predicted trend of buckling with varying salt has been observed experimentally [11]. The total length of DNA for 10 mM and 100 mM experimental data sets [11] differ slightly (*∼* 0.2 *μ*m); we renormalized the length of the 10 mM case (open squares) to be close to that of the 100 mM data (filled circles) for comparison in Figure 3a.

As the salt concentration is increased, the average number of superhelical turns per plectoneme domain increases, *i.e.*, the average size of each plectoneme domain increases. This effect is directly related to the decrease in DNA excluded volume at higher salt concentrations (Table I), which stabilizes the superhelical bends in a braid plectoneme.

In the supercoiled single DNA case, the opposite trend is observed, where lowering the salt concentration of the solution makes plectonemic buckling occur at a higher supercoiling density [47]. In stretched supercoiled DNA,the stability of the force-extended phase is unaltered by changing the ionic strength, but the supercoiled plectoneme phase (containing intra-molecular writhes) is relatively destabilized on decreasing the salt concentration, again due to the increase in effective diameter of the double helix, resulting in the observed trend of buckling at a lower linking number for higher salt concentrations.

The torque in the braid plotted as a function of the catenation (Figure 3b) shows a non-linear increase and a small non-monotonic overshoot at the buckling transition at all salt conditions. For a given catenation, the torque is higher for lower salt concentrations due to effective “swelling” of the braid. The torsional stress in the braid decreases with increasing salt, a trend also observed in supercoiled DNAs [42]. At 2 pN, we find nucleation of multiple domains of plectonemes at lower salt concentrations (*<*500 mM, see Figure 3c), while single plectoneme domain is favored at higher salt concentrations (*≈*500 mM). At lower salts, larger braid diameter increases the energy associated with superhelical bending of the braid in the plectoneme, making the formation of looped structures of braid favored over a superhelical structure, consequently favoring formation of multiple domains of plectoneme. We also find that the multiple domain structure of the plectoneme is favored at higher forces, *e.g.*, at 500 mM salt and forces *>* 3 pN more than one domain of plectoneme is energetically favorable. Post buckling, nucleation of plectoneme domains per added linking number is similar at lower salt concentrations, seen as the constant slopes in 10, 50 and 100 mM cases in Figure 3c. The length of the straight phase helical section increases faster for lower salt conditions due to the larger radius of the braid. The total size of the helical wrappings in the straight phase peaks at the buckling transition, after which double helix length is transferred into the growing plectoneme phase (Figure 3d).

At high salt concentrations (*>* 500 mM), we find a reversal of the above-mentioned trend with salt, *i.e.*, buckling transition occurs at a lower catenation for increasing salt (Figure 8). It may be possible to experimentally observe this effect, although we caution that at such high salt concentrations the applicability of the Debye-Hückel theory is questionable at best.

**Fig. 8.**
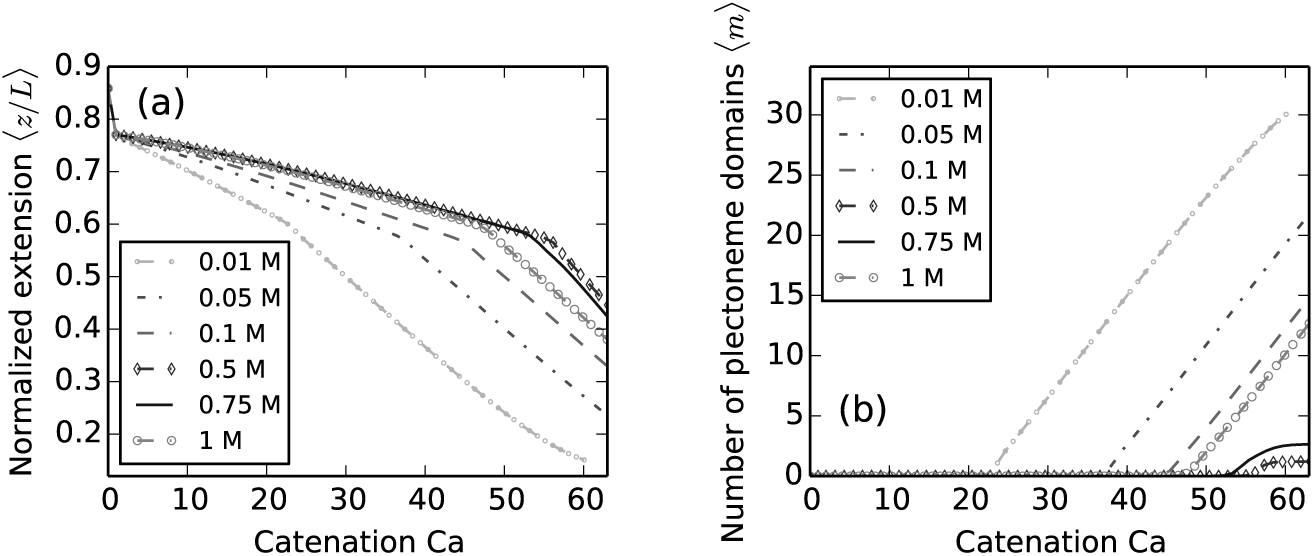
(a) Extension versus catenation for braids with *L* = 3.6 *μ*m, *d* = 0.42*L*, and *f* = 2 pN external force under a range of salt concentrations. Note that the buckling transition point is delayed on increasing the salt concentration till 0.5 M, after which the transition occurs at a lower catenation. (b) Number of plectoneme domains versus catenation for various salt concentrations. The multiple-domain character of the plectoneme phase also shows a non-monotonic trend: multiple domains are favored at very high salt concentrations *>* 500 mM. However, at such high salts we are in an extreme limit of the Debye-Hückel theory.

### C. Effect of the intertether distance

The intertether distance *d* between the two braided molecules affects the critical catenation, *i.e.,* the catenation at which buckling occurs (Figure 4a). Braiding molecules with larger *d* makes a helix with a larger aspect ratio (ratio of radius to pitch of the helix), which causes a steeper increase of the torque in the braid (Figure 4b). In effect, the twist modulus of the braid is larger for larger intertether distances and causes buckling at a lower catenation (Figure 4a). The torque at which a braid buckles is a thermodynamic property dependent on the external force and remains roughly the same on changing the intertether distance (Figure 4b) or the salt conditions (Figure 3b).

As mentioned before, the difference in extension between the states Ca = 0 and 1 is dependent on the intertether distance, but the two intertether distances at the two ends of the braid need not be the same. In fact, these distances being uncontrolled parameters in experiments, are almost never equal. Now, the theoretically predicted extension plots are a characteristic of the arithmetic mean of the intertether distances at the two ends of the braid; although, not surprisingly, the structure of the braid depends on the specific values of the two intertether distances (Figure 9a). In the case of unequal intertether distances, we find that the end-region associated with the larger intertether distance is bigger in size, *i.e.*, the helically wrapped region of the braid does not form at the center of the structure but is pushed towards the end with the smaller intertether distance.

**Fig. 9.**
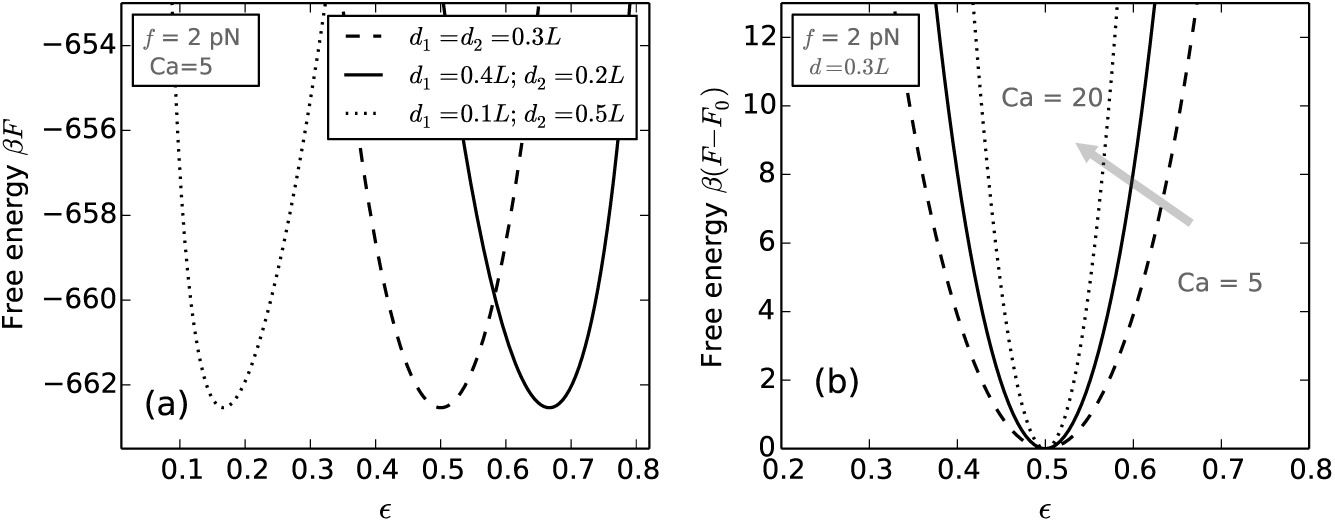
Effect of the intertether distance. (a) Comparison of the free energy versus *E* (Eq. D.1) for braids with various choices of the two intertether distances *d*1 and *d*2, but keeping the arithmetic mean of the two distances the same. The end-region where the two DNAs are closer to each other is smaller in size (*i.e.*, has less DNA in it), and vice-versa. (b) Free energy versus *E* for braids at various catenations (Ca=5, 12 and 20) with identical intertether distances *d* = 0.3*L*. The stiffness of the free energy decreases for lower catenation values indicating higher fluctuations in the relative sizes of the two end regions. The free energy curves have been shifted in the y-axis in order to overlay them.

We also find that the energy cost of fluctuations in the relative size of the two end regions, *i.e.*, small displacement of the entire helical section away from the equilibrium position are *O*(*kBT*), hence permissible, especially in the regime of low catenation (Figure 9b). The energy cost increases with increasing catenation, reflecting the sliding of the helical section is energetically expensive when the torque in the braid is higher. Such behavior of a braid may be possible to probe in braiding experiments done on DNA molecules labeled along their length with fluorescent tags.

### D. Braiding short DNA molecules

Braiding DNA molecules shorter than *≈*3 kb in size shows the same qualitative trends of extension decrease with catenation and formation of multiple buckled domains past the critical catenation density, as seen for larger molecules. However, due to the small size of the molecules and hence the smaller number of fluctuation states, discrete nucleation of buckled domains may be observed as steps in the extension plot (Figure 5a). The torque also shows multiple overshoots associated with nucleation of buckled domains (Figure 5b and 5c). Similarly, a step-like behavior can be seen in the length transfer between the straight and the plectoneme phase. The discrete jumps in extension or torque are masked by thermal fluctuations in longer molecules (4 kb or 1.3 *μ*m). Similarly, at lower forces, large fluctuations make the average extension of the braid decrease more uniformly (Figure 5a).

**Fig. 5.**
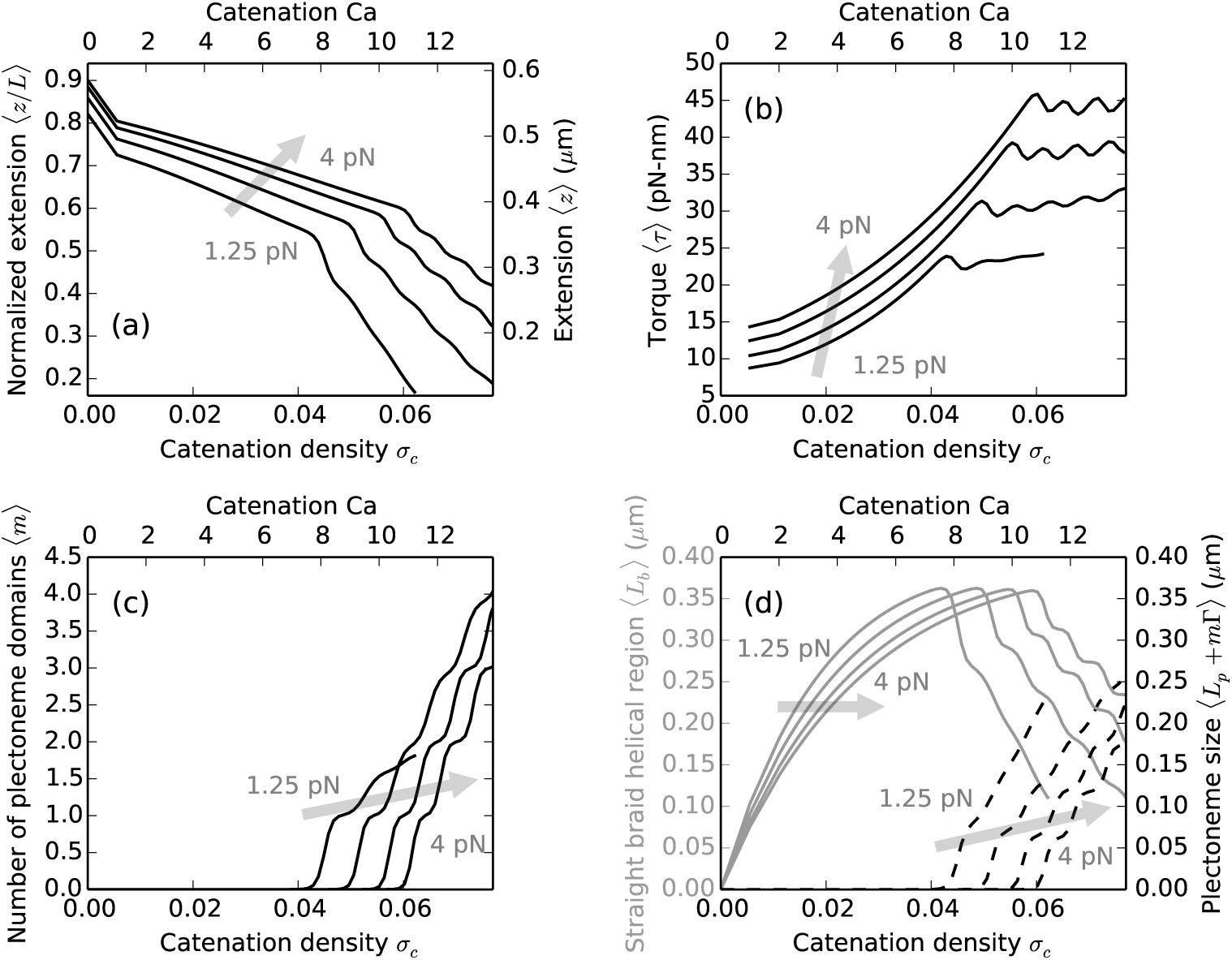
Braiding *≈*2 kb (*L* = 0.65 *μ*m) DNA at 100 mM Na+ with forces *f* = 1.25, 2, 3 and 4 pN, where the shaded arrows show the direction of increasing force. The intertether distance is 0.26 *μ*m (*d* = 0.4*L*), where the top and the bottom x-axes show catenation and catenation density (*σ*_*c*_ = Ca/Lk_0_) in the braid. (a) Relative extension (left y-axis) or extension (right y-axis) versus catenation for short DNA molecules. (b) Torque vs catenation shows multiple discrete “overshoots”, corresponding to the nucleation of new plectoneme domains. (c) Number of plectoneme domains vs catenation. The appearance of new plectoneme domains coincides with the steps in the extension or overshoots in the torque. (d) The size of the straight-phase helical region 〈*L*_*b*_〉 (left y-axis, solid gray curves) and the size of the plectoneme phase (*L*_*p*_ + *m*Γ)〉 (right y-axis, dashed black curves) versus catenation, where steps are associated with the formation of finite-length braid end loops. The successive nucleation events are smoothed out by thermal fluctuations in braids made up of long DNA molecules (≳4 kb).

## IV. CONCLUSION

We have presented a statistical-mechanical theory for the mechanics of a DNA braid, or a pair of catenated DNAs, as a function of catenation number and applied force (Figure 2). Our results are in good quantitative agreement with available data, and make a number of predictions for future experiments, including the nonlinear nature of the torque associated with DNA interwinding. A novel feature of DNA braids is a twist-dependent twist rigidity (Figure 2b) that arises from the lack of an intrinsic elastic twist stiffness for two catenated DNAs.

For the first time, we have quantitatively described the onset of the post-buckled state of the braid, and we find that the post-buckled “supercoiled” state is actually composed of many small buckled structures, each terminated by a small “end loop” (Figure 2c). This situation is reminiscent of the many-plectoneme state for a supercoiled single DNA at low salt [27, 28, 53]; however, for braids, this multi-buckled-structure state occurs for a broader range of salt concentrations (*<*0.5 M, see Figure 3c). The reason for the prevalence of the multi-buckled state is simply that the braided DNAs are relatively bulky, mimicking the effect of the larger excluded diameter that drives the multi-plectoneme state for single twisted DNAs. Inside the braid, there are electrostatic interactions between the two DNAs that force it to increase in radius, driving an increase in effective braid twist modulus as salt concentration is reduced (Figure 3b). This gives rise to a decrease of catenation number required for the onset of buckling as the salt concentration is decreased (Figure 3), the opposite of the trend observed for individual twisted double-helix DNAs. For the same reason, braid extension at fixed catenation number and force decreases with decreasing salt concentration (Figure 3a), again distinct from what is observed for twisted double-helix DNAs.

Under physiological conditions (100 mM salt), past the buckling transition point, the proliferation of new buckled domains leads to a decrease in the average size of each domain, *i.e.*, the average number of superhelical turns in a plectoneme domain decreases as the catenation is increased in the buckled braid. Now, an increase in the salt concentration, followed by a decrease in the DNA excluded volume (Table I) results in a decrease of the curvature energy associated with the superhelical bends in a plectoneme. Consequently, at high salt (≈0.5 M), the average number of superhelical turns per domain increases past the buckling point, leading to the formation of long superhelical structures instead of the appearance of multiple domains (Figure 3).

We also find that the mechanics of the braid and its buckling behavior are sensitive to the distance between the tethered DNA ends (Figure 4). Increased intertether distance leads to lower extensions (Figure 4a), higher torques, and buckling at lower catenation number (Figure 4b). We note that as the intertether distance is changed at a fixed force, the torque at the buckling point is nearly constant, *e.g.*, roughly 30 pN·nm at a force of 2 pN (Figure 4b).

The nucleation of the braid end loop at the buckling transition can likely be observed experimentally, possibly as a discontinuity in the extension versus catenation (Figure 2a) or dynamic switching of extension or as an overshoot in the torque versus catenation plots (Figure 2b). When the torsional stress in the braid is close to the critical buckling torque, the buckled state, separated by the nucleation energy cost becomes thermally accessible, resulting in equilibrium fluctuations between the straight and the buckled phase. Experimental studies [46, 47] on supercoiled single DNAs under torsional constraint have observed the discrete nucleation of the first plectoneme domain, which is associated with the abrupt nucleation of the plectoneme end loop. This abrupt transition has also been analyzed theoretically [26–28, 54], discussing the trends of the magnitude of the extension discontinuity with varied external force, salt concentration and contour length of the duplex DNA.

This paper qualitatively improves the way that fluctuations of the duplex DNAs in the braid around their average helical shape are accounted for. Instead of the semiquantitative scaling-like approach taken in prior works [20, 21, 28, 30, 32, 39], we systematically analyze the fluctuations of the DNAs around their average helical shape; we find that these fluctuations are small enough that it makes sense to think of the conformation of the braid as a helix (Figure 7), rather than the more disordered structure implicitly assumed in prior works.

As mentioned before, we assume spatially-uniform mean-field curvature in the braiding helices, and the introduction of a space-varying component to the helical curvature will be an interesting addition to the model. The effect of the spatially-varying helical pitch has been studied for loaded plies at zero temperature [34–36] as well as for straight DNA braids [30, 31]. Variable pitch solutions for many persistence-lengths long braided helices feature a constant helical angle inside the braid, which is smaller than the end angle that connects the helices with the end regions [30]. We note that we determine the end angle *φ* from free energy minimization, and we indeed find that an end angle larger than the helical angle *δ* is energetically favored, consistent with the effect observed in Ref. [30].

It may be interesting to study short braided DNAs (*≈*2 kb or 650 nm) because we find that for molecules of that short length, the successive addition of small buckled domains leads to a series of buckling transitions (Figure 5), possibly observable as steps in the extension. This is directly related to less thermal fluctuations in smaller DNA molecules. The effect of variable curvature is more prominent in short braids, which may provide additional fluctuations that further mask discrete jumps in extension for short DNA braids.

We also present a comprehensive treatment of the electrostatic effects (Appendix A) in helical braids, which will be useful for other problems where two DNAs are wrapped around one another.

## ACKNOWLEDGMENTS

The authors would like to acknowledge support from the National Science Foundation (Grants DMR-1206868 and MCB-1022117), the National Institutes of Health (Grant R01-GM105847 and U54-CA193419 [CR-PS-OC], and by subcontract to the University of Massachusetts under Grant U54DK107980 [Center for 3D Structure and Physics of the Genome]), and the Molecular Biophysics Training Program at Northwestern University.

## Appendix A Self-electrostatic energy of DNA braids

The minimum energy state of a charged rod in an ionic solution is the straight configuration, such that the electrostatic repulsion between the length elements of the rod is minimized. External work has to be done against the electrostatic repulsion to keep the rod bent in a helix shape, which is stored as the self energy. Prior theoretical works on braids [6, 30] as well as supercoiled single DNA plectonemes [20, 27, 28, 39] ignore the Debye-Hückel self energy of helically intertwined DNAs. In the following, we numerically compute the self energy in braids and propose an empirical fit function. We find that the energy contribution from Debye-Hückel-type self-interaction of the coaxial helices increases for helices with higher aspect ratio (ratio of radius to pitch), and is non-negligible for braids under 2 pN force at 100 mM monovalent salt concentration (Figure 6c and 6d).

We parameterize the helical arc length by a rotation angle *u* such that the distance *ρ* between any points on the helix is given by,

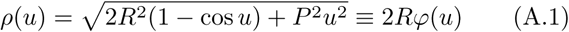

where *R* and 2*π P* are radius and pitch of the helix respectively; we define 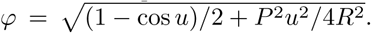. The authors of Ref. [39] used a similar parameterization scheme to study the enhancement of electrostatic interaction energy between two helically intertwined strands of a plectoneme as a function of the helix angle.

We write Ψ_*s*_, the total self-electrostatic potential for a braid of length *L* by integrating the spherically symmetric solution of the Debye-Hückel equation:

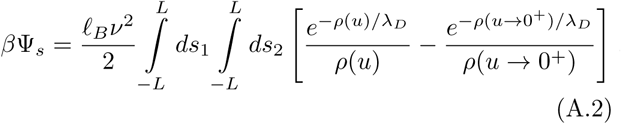

Now, the self-electrostatic potential per unit length *A* of the braid is given by *𝒰*_*s*_ ≡ Ψ_*s*_/(*L/A*), such that

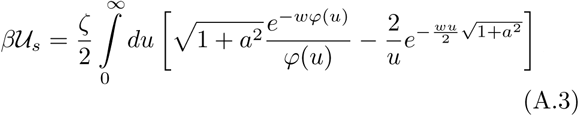

where *a ≡ P/R*, is the inverse of the aspect ratio of the helix, and *w ≡* 2*R*/*λ*_*D*_, is the scaled diameter of the helix. We have defined *ζ* = 2*A*ℓ_*B*_*v*^2^, where *ℓ*_*B*_ is the Bjerrum length and *?* is the effective linear change density in inverse-length units (Eq. 6 and Table I). We have shifted the reference of the free energy to subtract the contribution of the self-electrostatic energy of a straight rod.

We numerically evaluate the self-energy integral (Eq. A.3) for a range of practically relevant values of *w* and *a*, and propose the following empirical function that captures the behavior of the self-energy functional,

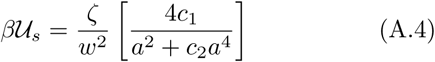

Figure 6a and 6b shows the comparison between the numerically evaluated value of the integral (Eq. A.3) and the empirical fit-function (Eq. A.4) with *ζ* = 1, *c*_1_ = 0.042 and *c*_2_ = 0.312.

Finally, we write the electrostatic potential energy per unit length *A* of braid with radius *R* and helix angle *δ*:

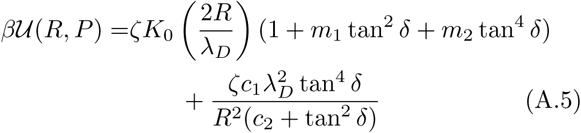

The first term is the interaction potential with *m*1 = 0.828, *m*2 = 0.864 [39], and the second term is the self-energy contribution, where *c*1 = 0.042 and *c*_2_ = 0.312.

Figure 6c and 6d shows the comparison between the self and the interaction energy component of the total electrostatic potential for various braid radii (*R*) and braiding angles (*δ*). For typical values of the braiding angle *≈25°* (Table I), the self-energy contribution becomes significant in braids with radius ≳ 3 nm, which corresponds to to ≳2 pN stretching tension on the braid at 100 mM Na+ (Figure 7a).

## Appendix B Radial uctuations

The average energy corresponding to radial fluctuations for each wavenumber *q* can be obtained from the fluctuation Hamiltonian (Eq. 9),

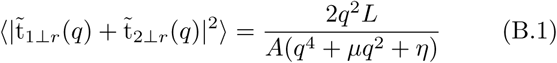

Now, the two-point correlation function associated with radial undulations can be computed as follows,

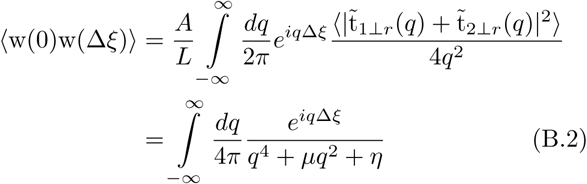

where Δ*ξ* is the distance between the two points. Note, the two-point correlation of radial fluctuations decays exponentially with the distance between the points: 〈(*w*(0)*w*(Δ*ξ*)〉 ∼ exp(*−k*Δ*ξ*), where 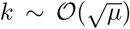, which is typical of Gaussian fluctuations.

We obtain the radial fluctuations in the braid from the zero-distance behavior of the above correlation function:

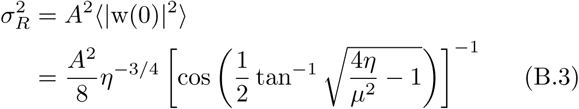

where *σ*_*R*_ is the fluctuation in braid radius. The radial probability distribution:

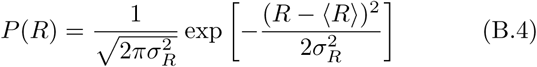

Figure 7a shows the probability distribution of braid radius for various forces. Higher forces result in a smaller average radius of the braid with less radial fluctuations. Figure 7b shows the variation of the mean and the fluctuation in braid radius for various forces at 100 mM salt. Radial fluctuations decrease with increasing force and are much smaller than the average value of the radius, suggesting small fluctuations of the braiding strands about their average shape.

## Appendix C Braids at high salt concentration

We find that the position of the buckling transition varies non-monotonically with the concentration of univalent salt in the solution. The amount of catenation at which the braid buckles increases with the salt concentration in the range 0.01 to 0.5 M, while decreases for higher salts (*>*0.5 M) (Figure 8a). We use *v* = (47.8, 78.1) nm^−1^ corresponding to (0.75, 1) M salt concentrations respectively [39]. We also find that for 2 pN force at salt concentrations *≈*0.5 M single domain plectoneme is favored over multiple domains, but multiple domains are again favored for *>* 0.5 M salt (Figure 8b). However, at salt concentrations above a few hundred millimolars, the Debye length of the solution is smaller than the DNA radius (Table I), which is an extreme limit of the mean-field Debye-Hückel theory.

## Appendix D Intertether distance

For a braid with unequal intertether distances *d*1 and *d*_2_, we define a dimensionless parameter *∊* such that

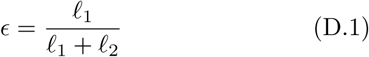

where the total length in the end-regions *L*_*e*_ (Eq. 15) is divided into *ℓ*_1_ and *ℓ*_2_ corresponding to the two endregions. Hence, *c* = 1*/*2 indicates the scenario of a braid with symmetric end-regions. Figure 9a shows the minimized total free energy (Eq. 15 and 16) versus *c* for three pairs of intertether distances, keeping the arithmetic mean of the distances the same in all three choices. The extension plots are a characteristic of the arithmetic mean, but the relative size of the end-regions depends on the particular choice of *d*_1_ and *d*_2_. The end-region associated with the larger of the two intertether distances contains more length of double-helix DNA.

Figure 9b shows minimized free energy versus *∊* for a braid with symmetric ends but various catenations. The stiffness of the potential near the equilibrium value of *∊* increases with increasing catenation in the braid. This suggests the possibility of fluctuation in the relative sizes of the end-regions, which would be higher for lower catenations and becomes energetically expensive in the regime of tight braid or braid with high catenation density. It may be possible to directly visualize the sliding of the braid helical intertwines in a braiding experiment using DNA molecules labeled with fluorescent tags.

